# Optimal control of acute myeloid leukaemia

**DOI:** 10.1101/429811

**Authors:** Jesse A Sharp, Alexander P Browning, Tarunendu Mapder, Kevin Burrage, Matthew J Simpson

## Abstract

Acute myeloid leukaemia (AML) is a blood cancer affecting haematopoietic stem cells. AML is routinely treated with chemotherapy, and so it is of great interest to develop optimal chemotherapy treatment strategies. In this work, we incorporate an immune response into a stem cell model of AML, since we find that previous models lacking an immune response are inappropriate for deriving optimal control strategies. Using optimal control theory, we produce continuous controls and bang-bang controls, corresponding to a range of objectives and parameter choices. Through example calculations, we provide a practical approach to applying optimal control using Pontryagin’s Maximum Principle. In particular, we describe and explore factors that have a profound influence on numerical convergence. We find that the convergence behaviour is sensitive to the method of control updating, the nature of the control, and to the relative weighting of terms in the objective function. All codes we use to implement optimal control are made available.

## 1 Introduction

Acute Myeloid Leukaemia (AML) is a blood cancer that is characterised by haematopoietic stem cells, primarily in the bone marrow, transforming into leukaemic blast cells [22,47]. These blast cells no longer undergo normal differentiation or maturation and stop responding to normal regulators of proliferation [23]; their presence in the bone marrow niche disrupts normal haematopoiesis [22]. AML has significant mortality rates, with a five-year survival rate of 24.5% [8], and challenges in treatment arise not only in eradication of the leukaemic cells but also prophylaxis and treatment of numerous life threatening complications that arise due to the absence of sufficient healthy blood cells [62]. Multiple interventions are employed in the management and treatment of AML, including: leukapheresis; haematopoietic stem cell transplants; radiotherapy; chemotherapy and immunotherapy [4,47,52].

Mathematical models are widely used to gain insight into complex biological processes [29,48]. Mathematical models facilitate the development of novel hypotheses, allow us to test assumptions, improve our understanding of biological interactions, interpret experimental data and assist in generating parameter estimates. Furthermore, mathematical models provide a convenient, low-cost mechanism for investigating biological processes and interventions for which experimental data may be scarce, cost-prohibitive or difficult to obtain owing to ethical issues. Mathematical models are routinely used to interrogate a variety of processes relating to cancer research including: incidence; development and metastasis; tumour growth; immune reaction and treatment [13,16,22,31,43,59]. Recently, mathematical models have been used to investigate various aspects of AML, including: incidence [41]; pathogenesis [19]; interactions between cancer and healthy haematopoietic stem cells within the bone marrow niche [22]; and recurrence following remission [50].

Determining how to apply optimally a treatment such as chemotherapy is of great practical and theoretical interest. Chemotherapy, a common treatment for AML [21], is associated with significant health costs related to the cytotoxicity of chemotherapeutic agents [11,47], but also substantial economic cost [64]. Optimal control theory provides us with tools for determining the optimal way to apply a control to a model such that some desired quantities of interest are minimised or maximised. Further, it facilitates assessment of the efficacy of hypothetical treatment protocols relative to a theoretical optimal treatment. Optimal control has been applied to a range of medically motivated biological models recently; including vaccination, tumour therapy and drug scheduling [15,17,35,36,44].

In this work we consider a recent haematopoietic stem cell model of AML [22]. After examining the steady state behaviour associated with this model, we make a biologically appropriate and mathematically convenient modification by incorporating an immune response in the form of a Michaelis-Menten kinetic function. Overall, in this work we pursue two broad aims:

1. Determine how to apply optimal control to the model, accounting for key clinical features such as the competition between the negative effects of the disease and the negative effects of the treatment;
2. Provide a concise and insightful discussion of the methodology and numerical implementation of optimal control, as we find that much of the existing literature is opaque with regard to practical implementation.

In addressing these aims, we provide a brief introduction to the theory of optimal control and apply optimal control techniques to the modified model, identifying optimal treatment strategies under a variety of circumstances. This leads us to consider both continuous and discontinuous bang-bang optimal controls. Our work provides a comprehensive discussion of practical issues that can arise when applying optimal control, and we explore key factors that influence numerical convergence when using a forward-backward sweep algorithm to solve two-point boundary value problems that arise. The code we use to implement the algorithms associated with the optimal control solutions is freely available on GitHub.

In Section 2 we present a haemotopoietic stem cell model of AML [22], and discuss the steady states. In Section 3 the importance of an immune response is outlined, and the model is modified to include such a response. In Section 4, we present discussion and results of optimal control applied to the modified AML model. Finally, concluding remarks are provided in Section 5. In the supplementary material document we extend the work in this document to consider: (i) arbitrary initial conditions, and; (ii) controls that impact multiple species.

## 2 Acute myeloid leukaemia model

Crowell, MacLean and Stumpf [22] propose a system of ordinary differential equations (ODEs) to model AML. Their model can be written as,

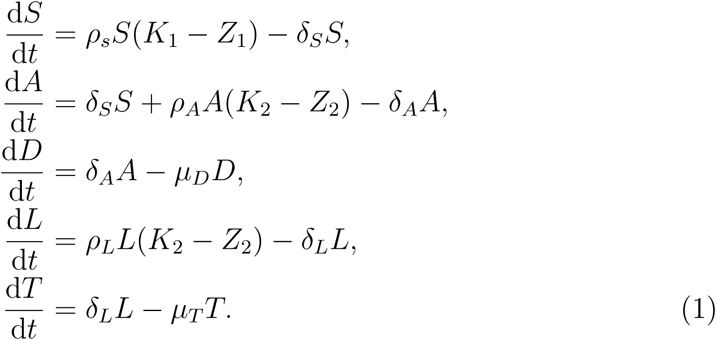

Here *S*(*t*), *A*(*t*), *D*(*t*), *L*(*t*) and *T* (*t*) represent haematopoietic stem cells, progenitor cells, terminally differentiated cells of *S*(*t*), leukaemia stem cells and fully differentiated leukaemia cells, respectively. *Z*_1_(*t*) = *S*(*t*) and *Z*_2_(*t*) = *A*(*t*) + *L*(*t*), where *A*(*t*) and *L*(*t*) are coupled as the proliferating leukaemia population (*L*(*t*)) competes with the haematopoietic progenitor cell population (*A*(*t*)). This competition is motivated in [22] by the hypothesis that leukaemic stem cells and haematopoietic stem cells occupy the same niche within the bone marrow [26,58] and hence compete for resources. This niche interaction has been demonstrated as being crucial to similar haematopoietic and leukaemic cell models of chronic myeloid leukaemia [43]. Throughout this work we present numerical solutions to this model and other related models. In all solutions presented the parameters are dimensionless, such that the time scale is arbitrary and cell population sizes within the bone marrow are expressed as a portion of the carrying capacities; *K*_1_ = *K*_2_ = 1. Setting these carrying capacities to be of equal size is a simplifying assumption in our analysis, though we note that this is not required, and could be relaxed if suitable alternative estimates of the carrying capacities were identified.

Crowell, Mac Lean and Stumpf use numerical solutions of Equation (1) to identify parameter values that lead to particular long time steady state solutions of the model. In this work we will use standard variables to denote time dependent quantities, such as *S*(*t*), and an overbar to denote long-time steady quantities, such as 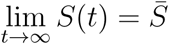. The parameters we use are summarised in Table 1, and we note that the model supports three non-trivial steady states:

**Table 1:** Parameters values used in this work.

| Parameter description | Value |
| --- | --- |
| Proliferation of $S$ | $\rho_S = 0.5$ |
| Proliferation of $A$ | $\rho_A = 0.43$ |
| Proliferation of $L$ | $\rho_L = 0.27$ |
| Differentiation of $S$ into $A$ | $\delta_S = 0.14$ |
| Differentiation of $A$ into $D$ | $\delta_A = 0.44$ |
| Differentiation of $L$ into $T$ | $\delta_L = 0.05$ |
| Migration of $D$ into the blood stream | $\mu_D = 0.275$ |
| Migration of $T$ into the blood stream | $\mu_T = 0.3$ |
| Carrying capacity of the compartment with $S$ | $K_1 = 1$ |
| Carrying capacity of the compartment with $A$ and $L$ | $K_2 = 1$ |
| Characteristic rate of the immune response | $\alpha = 0.015$ |
| Half saturation constant of the immune response | $\gamma = 0.01$ |

1. The *coexisting* steady state requires 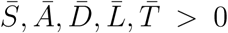 simultaneously. In this work we are interested in modelling the optimal application of an intervention (or control) such as chemotherapy to the system that shifts it from the coexisting steady state towards the healthy steady state. Examples trajectories resulting in the coexisting steady state are given in Figure 2a and Figure 2b.
2. The *healthy* steady state consists of 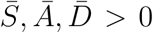 and *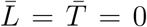*, such that there is a population of each healthy cell species and no leukaemia is present. The healthy steady state is demonstrated in Figure 2c.
3. The third steady state is *leukaemic*, characterised by 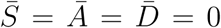 and 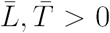, such that only leukaemic cells are present. The leukaemic steady state is demonstrated in Figure 2d.

The leukaemic steady state is less interesting from an intervention perspective as it cannot be steered towards the healthy steady state via a control such as chemotherapy alone; requiring in addition a source of healthy cells.

Parameter values in Table 1 are used in all numerical solutions presented in this work, unless otherwise indicated. These values match those specified in [22] to produce a healthy steady state, noting that [22] included parameter sweeps over *ρ*_*S*_, *ρ*_*A*_, *δ*_*S*_ and *δ*_*A*_, with the exception of *δ*_*L*_. We have set *δ*_*L*_ = 0.05 to produce the coexisting steady state, although other values for *δ*_*L*_ also produce this coexisting steady state.

Schematics showing the key features of the original model, a modified model that incorporates an immune response (Section 3), and the modified model subject to a control (Section 4) are presented in Figure 1. Typical numerical solutions of the original model are presented in Figure 2. All numerical results presented in this study are obtained using a fourth-order Runge-Kutta method [53] with a constant time step of *δt* = 0.001. We find that this choice is sufficient to produce numerical solutions that are grid-independent. From the numerical results we observe that for the parameter values given in Table 1, provided that initially *S*(0) > 0 and *L*(0) > 0, the system will tend towards the coexisting steady state. In Section 3 we modify the model to incorporate an immune response, such that sufficiently small leukaemic populations will decay without intervention.

**Fig. 1.**
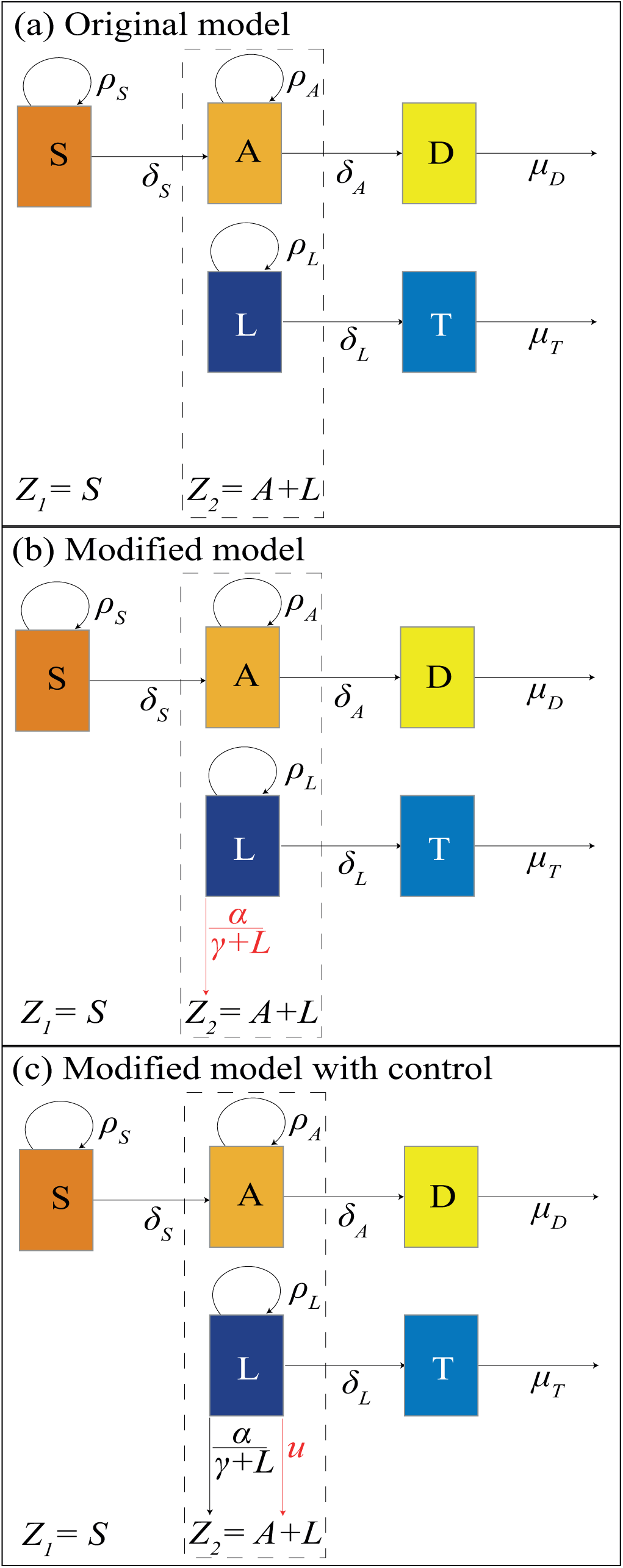
Schematics present the interactions and associated parameters for the (a) original model [22], (b) modified model with immune response and (c) modified model subject to a control, *u*. In each schematic the additional response is highlighted in red.

**Fig. 2.**
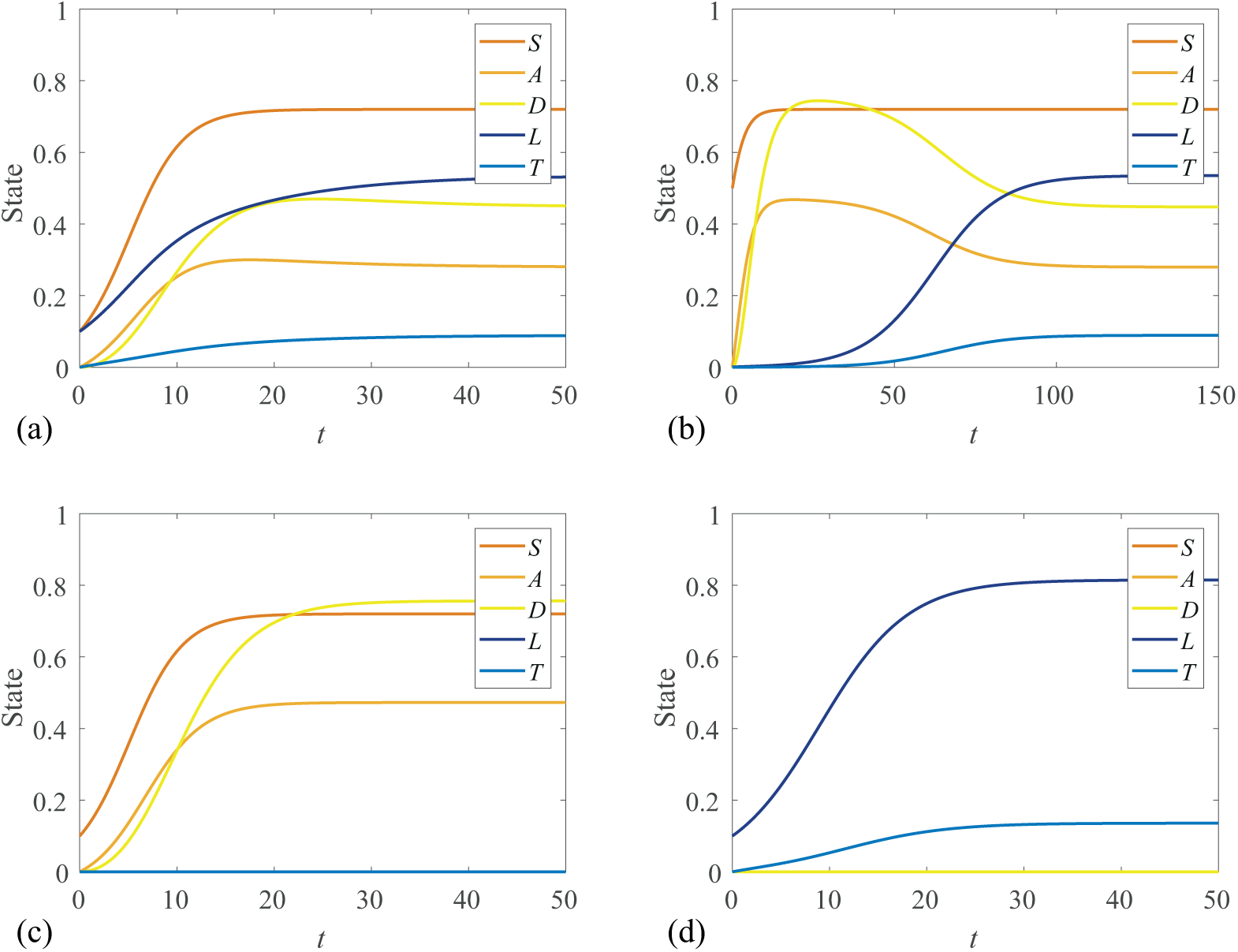
Numerical solutions of Equations 1 for various initial conditions: (a) Coexisting steady state solution with [*S*(0), *A*(0), *D*(0), *L*(0), *T* (0)] = [0.1, 0, 0, 0.1, 0]. (b) Coexisting steady state with [0.5, 0, 0, 10^−3^, 0]. (c) Healthy steady state with [0.1, 0, 0, 0, 0]. (d) Leukaemic steady state with [0, 0, 0, 0.1, 0].

In Figure 2b we note that although the initial leukaemia stem cell population is small compared to the initial haematopoietic stem cell population, the system eventually evolves to the same coexisting steady state as in Figure 2a. However, this steady state condition requires a longer timescale to develop from the different initial conditions.

## 3 Incorporating the immune response

The immune system is known to play a critical role in the development, metastasis, treatment and recurrence of cancers [25,27]. This knowledge is supported by a range of clinical evidence, including a well-documented increased risk of cancer incidence in patients with immunodeficiency [18]. This is exemplified by experimental mouse models where mice are typically immunocompromised to avoid transplanted cancers being destroyed by the immune response in xenograft studies [20]. Furthermore, tumours found in immunocompetent hosts are observed to exhibit mechanisms for avoiding immune response [46].

The behaviour exhibited in Figure 2b indicates that the system cannot reach a healthy non-leukaemic steady state in the presence of even small leukaemic stem cell populations. It is reasonable to expect that under some circumstances a small leukaemic population may be outcompeted by healthy cells occupying the same niche [42], without intervention. Therefore, we consider a modification to the model proposed by Crowell, MacLean and Stumpf to incorporate an immune response. We expect this immune response to be effective for small *L* and ineffective for large *L*, and so we mimic this by introducing a Michaelis-Menten term to represent the immune response, giving,

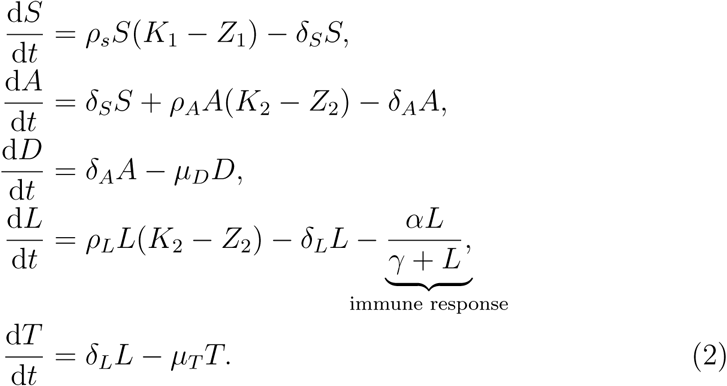

Including an immune response in the model is not only mathematically convenient in that it provides desirable steady states that we discuss later in this section, but also biologically relevant. Immune responses are widely studied in both the theoretical and experimental biology literature and acknowledged as an important contributor to pathogenesis and tumour dynamics in AML [7,32,61]. Additionally, immunotherapy is being investigated as an alternative to chemotherapy for treatment of AML and many other cancers [10,40,45].

Michaelis-Menten terms are commonly used to incorporate immune responses in other biologically motivated models [2,24,38]. However, it is unclear, simply by inspection, what parameter values are required to obtain two stable steady states: one coexisting and one healthy. For *γ* ≪ *α* the Michaelis-Menten term behaves as exponential decay at a rate of *α*, while for *γ* ≫ *L* it behaves as a linear sink term [56,57]. Intuitively, we expect setting *γ* = 𝒪 (*L*) will produce the desired dynamics whereby the immune response is effective for small *L* and ineffective for large *L*.

We investigate further by considering the potential steady states permitted by Equation (2). We note that *S* is governed by a logistic growth mechanism that does not depend on any of the other species so we have 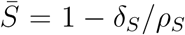. Similarly, *D* and *T* do not influence the other populations and hence can be neglected in the consideration of the steady states. Therefore, we consider a reduced system in terms of *A, L* with 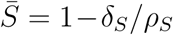, recalling that *Z*_2_ = *A*+*L*, and through scaling *K*_2_ = 1,

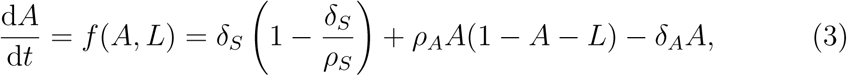

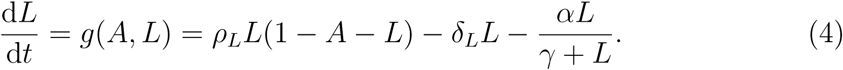

By inspection, there is a trivial L-nullcline at 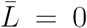. We can find the Anullcline by setting *f* (*A, L*) = 0 in Equation (3),

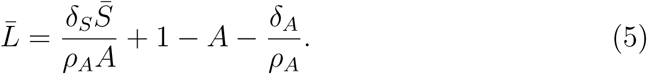

Similarly, we can find the non-trivial L-nullcline by setting *g*(*A, L*) = 0 in Equation (4),

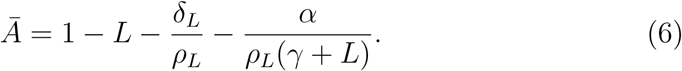

The nullclines, given by Equations (5) and (6), are hyperbolas. In Figure 3 we present phase planes for both the modified (with immune response) and unmodified (no immune response) models showing dynamics of the *A* and *L* populations within the physically meaningful region, *A* + *L* ≤ 1.

**Fig. 3.**
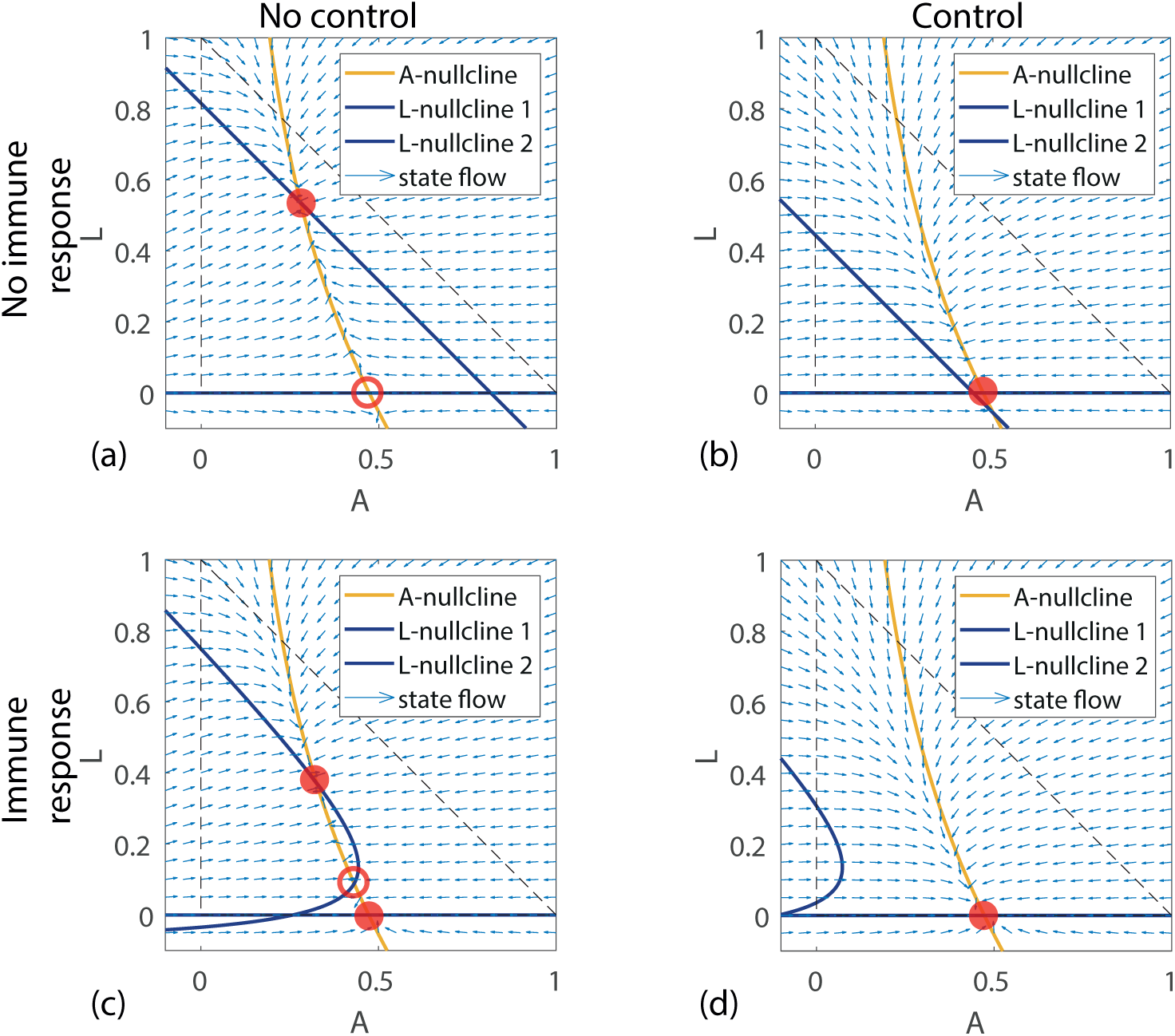
Nullclines and steady states of the model with (a) no immune response and no control, (b) no immune response with control, (c) immune response with no control and (d) immune response with control, using parameters for a coexistence steady state; [*ρ*_*S*_, *ρ*_*A*_, *ρ*_*L*_, *δ*_*S*_, *δ*_*A*_, *δ*_*L*_] = [0.5, 0.43, .027, 0.14, 0.44, 0.05]. Physically realistic fixed points are marked with closed discs if stable or open discs if unstable. The application of a control in (b) and (d) corresponds to *u* ≡ 0.1, effectively increasing *δ*_*L*_ to 0.15 (a control could be a treatment such as chemotherapy that increases the rate of decay of leukaemic stem cells, this is discussed in Section 4). In (c), for particular choices of the introduced parameters *α* and *γ* it is possible for the hyperbolas to intersect twice within the physically realistic region (dashed triangle). Figures (c) and (d) are produced with *α* = 0.015, *γ* = 0.1. Without an immune response, as illustrated in (a) and (b), application of a control can steer the system towards a stable healthy steady state, however this fixed point becomes unstable if the control is ceased, causing the system to revert to the coexisting steady state. With an immune response, as illustrated in (c) and (d), once the control steers the system into the attractor region of the healthy fixed point, the system does not revert to the coexisting steady state upon ceasing the control.

This system has the desired property that we outlined previously, namely that there is a stable steady state of coexistence that we aim to steer to the stable state with no leukaemia through applying optimal control. Numerical solutions of the modified model with no control are presented in Figure 4.

**Fig. 4.**
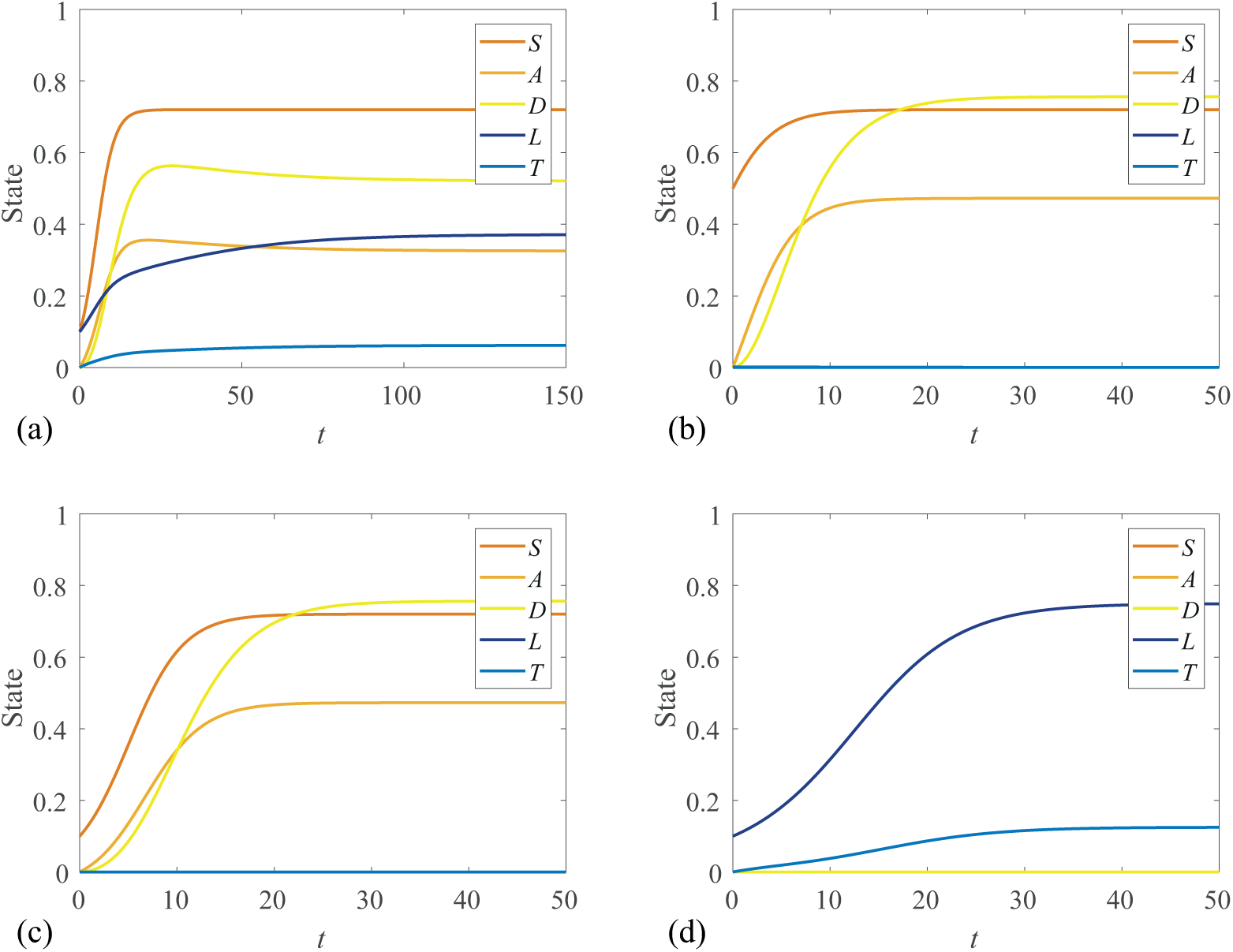
Numerical solutions to the modified model with an immune response for initial conditions corresponding to Figure 2. In (a) we observe coexistence, though it takes longer for the solutions to approach steady state when compared with the original model (Figure 2a). This result is presented over a larger time-scale. With the introduction of the Michaelis-Menten style immune response to leukaemia, we observe in (b) that a small leukaemia stem cell population does not survive in the presence of a haematopoietic stem cell population. This is in contrast to Figure 2b, where a minute population of leukaemic stem cells was sufficient to grow to a coexisting steady state. These figures are produced with immune response parameters *α* = 0.015, *γ* = 0.1.

## 4 Methods

In this section we provide a concise overview of the theory of optimal control. Methods for solving optimal control problems are discussed. We determine optimal controls to the model presented in Section 3. Specifically, we consider continuous optimal controls corresponding to quadratic pay-off functions and discontinuous bang-bang optimal controls corresponding to linear pay-off functions. Numerical solutions are produced for several different pay-off weighting parameter combinations.

### 4.1 Optimal control theory

The basic principle of optimal control is to apply an external force, the *control*, to a system of differential equations, the *state equations*, to cause the solution, the *state*, to follow a new trajectory and/or arrive at a different final state. The goal of optimal control is to select a particular control that maximises or minimises a chosen objective functional, the *pay-off*; typically a function of the state and the control. The pay-off is chosen such that the new trajectory/final state are preferred to that of the uncontrolled state, accounting for any cost associated with applying the control.

A typical optimal control problem will introduce the state equations as functions of the state **x**(*t*) and the control *u*(*t*), with initial state **x**(0) = **x**_**0**_,

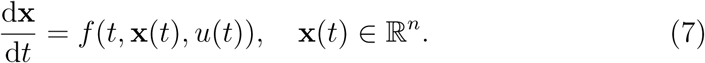

It is also necessary to specify either a final time *t*_*f*_ with the final state free, or a final state **x**(*t*_*f*_), with the final time free.

A pay-off function *J* is defined as a function of the final state, **x**(*t*_*f*_), and a cost function ℒ (*t,* **x**(*t*), *u*(*t*)) integrated from initial time (*t*_0_) to final time (*t*_*f*_). Through choosing an optimal control *u*\*(*t*) and solving for the corresponding optimal state **x**\*(*t*), we seek to maximise or minimise this objective function. Selecting the pay-off enables us to incorporate the context of our application and determine the meaning of *optimality*. In general, the pay-off function can be written as,

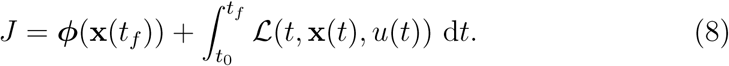

Depending on the form of ***ϕ***, it may be possible to incorporate ***ϕ*** into ℒ by restating the final state constraint in terms of an integral expression using the Fundamental Theorem of Calculus, and noting that ***ϕ***(**x**(*t*_0_)) is constant and hence does not impact the optimal control. The resulting unconstrained optimal control problem is often more straightforward to solve than the constrained problem.

The optimal control can be found by solving necessary conditions obtained through application of Pontryagin’s Maximum Principle (PMP) [51], or a necessary and sufficient condition by forming and solving the Hamilton-Jacobi-Bellman partial differential equation; a dynamic programming approach [9]. In this work we use the PMP and we construct the Hamiltonian, *H*(*t,* **x**, *u,* **λ**) = ℒ (*t,* **x**, *u*) + **λ***f,* where **λ** = [λ_1_(*t*), λ_2_(*t*), …, λ_*n*_(*t*)] are the adjoint variables for an *n*-dimensional state. The adjoint is analogous to Lagrange multipliers for unconstrained optimisation problems. Through the Hamiltonian, the adjoint allows us to link our state to our pay-off function. The necessary conditions can be expressed in terms of the Hamiltonian,

1. The optimality condition is obtained by minimising the Hamiltonian,

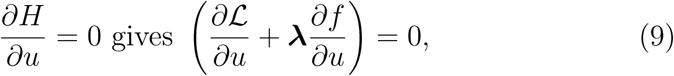
2. the adjoint, also referred to as *co-state*, is found by setting,

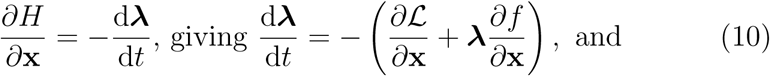
3. satisfying the transversality condition,

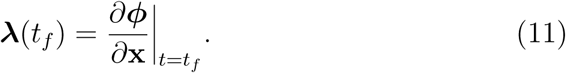

### 4.2 Continuous optimal control

In this section we consider optimal control applied to the AML model presented in Section 3. From this point we omit the implied time dependence of all control, state and co-state variables for notational convenience. Consider the steady states we observed for the coexistent parameter values of model 1. Suppose we wish to apply an optimal control that steers the system from a steady state observed in Figure 4a towards a healthy steady state (Figure 4b). This could be achieved by applying a drug *u*(*t*), the dosage of which may vary over time, that kills leukaemic stem cells,

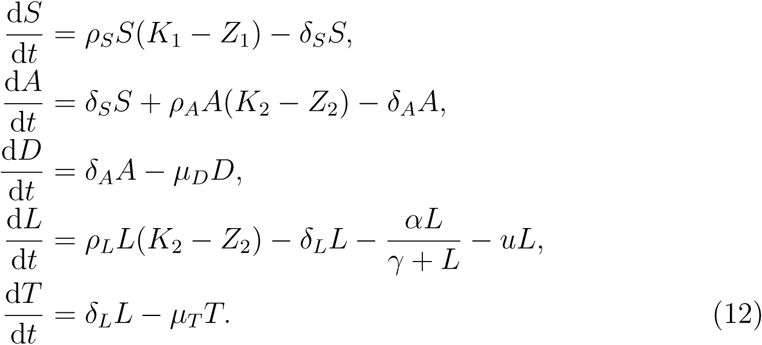

A potential pay-off function for this optimal control problem is to minimise,

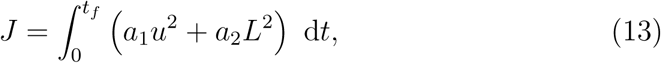

where the control problem is assumed to start at time zero and run until a fixed end time of *t*_*f*_. In defining a pay-off function there is significant scope for flexibility, and what constitutes an appropriate choice depends on the application. The parameters *a*_1_ > 0 and *a*_2_ > 0 are chosen to weight the importance of each term in the pay-off, and can be adjusted to best suit a particular application. Through scaling it can be seen that for this example only the relative weighting (*a*_1_*/a*_2_) is important, however we specify *a*_1_ and *a*_2_ separately for clarity.

Quadratic pay-off functions have several desirable mathematical properties that increase the ease of finding optimal solutions; they are smooth and have only a single extremum. Furthermore, quadratic pay-off functions help to avoid non-physical controls that may otherwise be found. For example; if the pay-off was a cubic function of *u*, setting *u* to be large and negative may minimise the pay-off but be physically unrealisable. Quadratic pay-off functions also have some desirable physical properties; a quadratic term will apply a harsher penalty to large amounts of control than small amounts [6], which in many treatments, such as chemotherapy, is desirable [30]. In control engineering applications, the control, *u*, is thought to be proportional to a voltage or current, in which case a quadratic pay-off has a convenient interpretation, as *u*^2^ is proportional to power, and the integral of this power over an interval is proportional to the energy expended [6]. Pay-off functions that are quadratic in the control variable are used in many biological [39,54] and engineering applications [3,49].

We can construct the Hamiltonian as *H* = ℒ+ **λ***f*; where *f* is the right hand side of Equation (12), **λ** = [λ_1_, λ_2_, λ_3_, λ_4_, λ_5_], and from Equation (13), we have ℒ=*a*_1_*u*^2^+*a*_2_*L*^2^, giving,

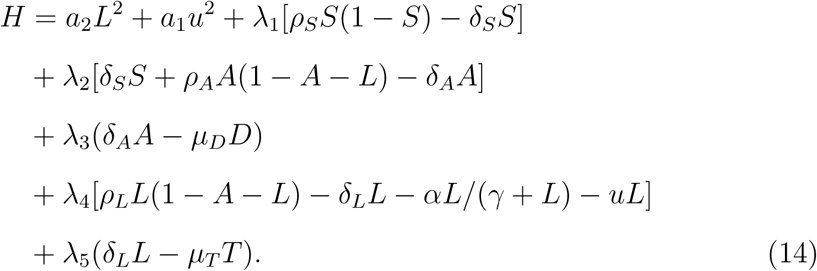

From Equation (9), we find the optimal control by setting *∂H/∂u* = 0, giving *u*\* = λ_4_*L/*2*a*_1_. Following Equation (10), the co-state equations for **λ** are found by setting d**λ***/*d*t* = −*∂H/∂***x**,

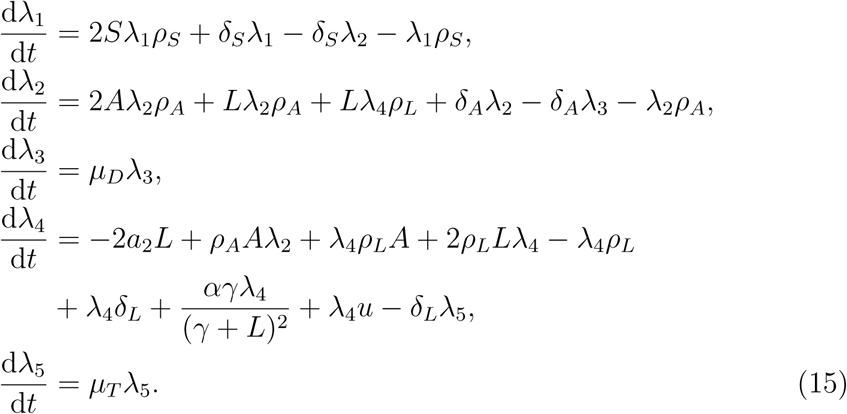

The transversality condition, Equation (11), gives final time conditions on the co-state, Equation (15); **λ**(*t*_*f*_) = [0, 0, 0, 0, 0]. Assuming that the initial state is known; [*S*(0), *A*(0), *D*(0), *L*(0), *T* (0)], it is now possible to determine the optimal control and corresponding state and co-state through solving a two-point boundary value problem (BVP).

We solve Equation (2) numerically to reach the stable coexistence steady state of the uncontrolled model. These steady state values in the absence of the control are used as the initial state conditions to solve the BVP to find the optimal control solution. The initial condition for the optimal control problem is [*S*(0), *A*(0), *D*(0), *L*(0), *T* (0)] = [0.7200, 0.3255, 0.5207, 0.3715, 0.0619]. Initialising the optimal control solution from the uncontrolled steady state is not necessary, however it helps to illustrate the role of the control. We demonstrate this flexibility by generating results for a range of arbitrary initial conditions and control start times. These results are presented and discussed in the supplementary material.

There are a range analytical methods available for solving some forms of BVP under certain conditions [1,63]. However, in this work we focus on numerical solutions with a view to identifying and discussing typical issues that may arise in implementation. Common numerical solution techniques include shooting and forward-backward sweep methods (FBSM) [28,37]. The most effective numerical method depends on the particular BVP. The single shooting method is relatively straightforward, but can be sensitive to the initial guess of the co-state. Forming a suitable guess for the initial values of the co-state is challenging, as the co-state does not have a straightforward physical interpretation. Although the FBSM calls for an initial guess for the control over the entire interval, this can often be straightforward to determine, as we will demonstrate.

We apply the FBSM using an initial guess for the control, *u*(*t*) ≡ 0, to solve for the state variables forward in time. The co-state is then solved backward in time. In each case a fixed step fourth order Runge-Kutta method is applied to solve the relevant system of ODEs. Using these solutions, the control is updated and the process is repeated until convergence is achieved. The algorithm for the FBSM is given in Algorithm 1.

#### Algorithm 1: Forward-backward sweep

i. Make an initial guess of *u*(*t*). *Typically u*(*t*) ≡ 0 *is sufficient, though a more thoughtful choice may result in fewer iterations required for convergence.*
ii. Using the initial condition **x**(0) = **x**_0_, solve for **x**(*t*) forward in time using the initial guess of *u*(*t*).
iii. Using the transversality condition **λ**(*t*_*f*_), solve for **λ**(*t*) backwards in time, using the values for *u*(*t*) and **x**(*t*).
iv. Calculate *u*_new_(*t*) by evaluating the expression for the optimal control *u*\*(*t*) using the updated **x**(*t*) and **λ**(*t*) values.
v. Update *u*(*t*) based on a combination of *u*_new_(*t*) and the previous *u*(*t*). *For continuous controls applied to relatively simple systems, it may be possible to use u*_*new*_(*t*) *directly* (*u*(*t*) = *u*_new_(*t*)), *however this is not sufficient to achieve convergence in general. We discuss this further in Section 4.4.*
vi. Check for convergence. *If* **x**(*t*), **λ**(*t*) *and u*(*t*) *are within a specified absolute or relative tolerance of the previous iteration, accept* **x**(*t*), **λ**(*t*) *and u*(*t*) *as having converged, otherwise return to Step* ii. *and repeat the process using the updated u*(*t*).

Solutions are provided in Figure 5 for various weighting on the control parameters. As expected, when *a*_1_ > *a*_2_, placing a greater weighting on the negative impact of the control than the negative impact of the leukaemic stem cells we observe that the control is applied at a lower level than when *a*_1_ < *a*_2_. When the pay-off weightings are equal, as shown in Figure 5b, the continuous control is applied at an amount similar to the level of the leukaemic stem cell population. Similarly, when the amount of control applied is larger, we observe that the leukaemic stem cell population declines at a faster rate. With *a*_1_ > *a*_2_, as in Figure 5c, we observe that the leukaemic population is effectively eradicated by *t*_*f*_, whereas when *a*_1_ < *a*_2_ we see, in Figure 5d, that a leukaemic population remains at *t*_*f*_. A limitation of specifying a fixed final time, as opposed to a fixed final state, is that the optimal outcome is dependent on the specified final time, and there is no consideration for what may happen *after t*_*f*_. In many applications, the notion of what happens beyond the control interval is not of interest, though in some instances specifying a final state may be more sensible. In this work we consider fixed final time problems for ease of comparison between controls under different parameter regimes, though we acknowledge that specifying a final state, such as *no leukaemic stem cells*, may be more biologically appropriate.

**Fig. 5.**
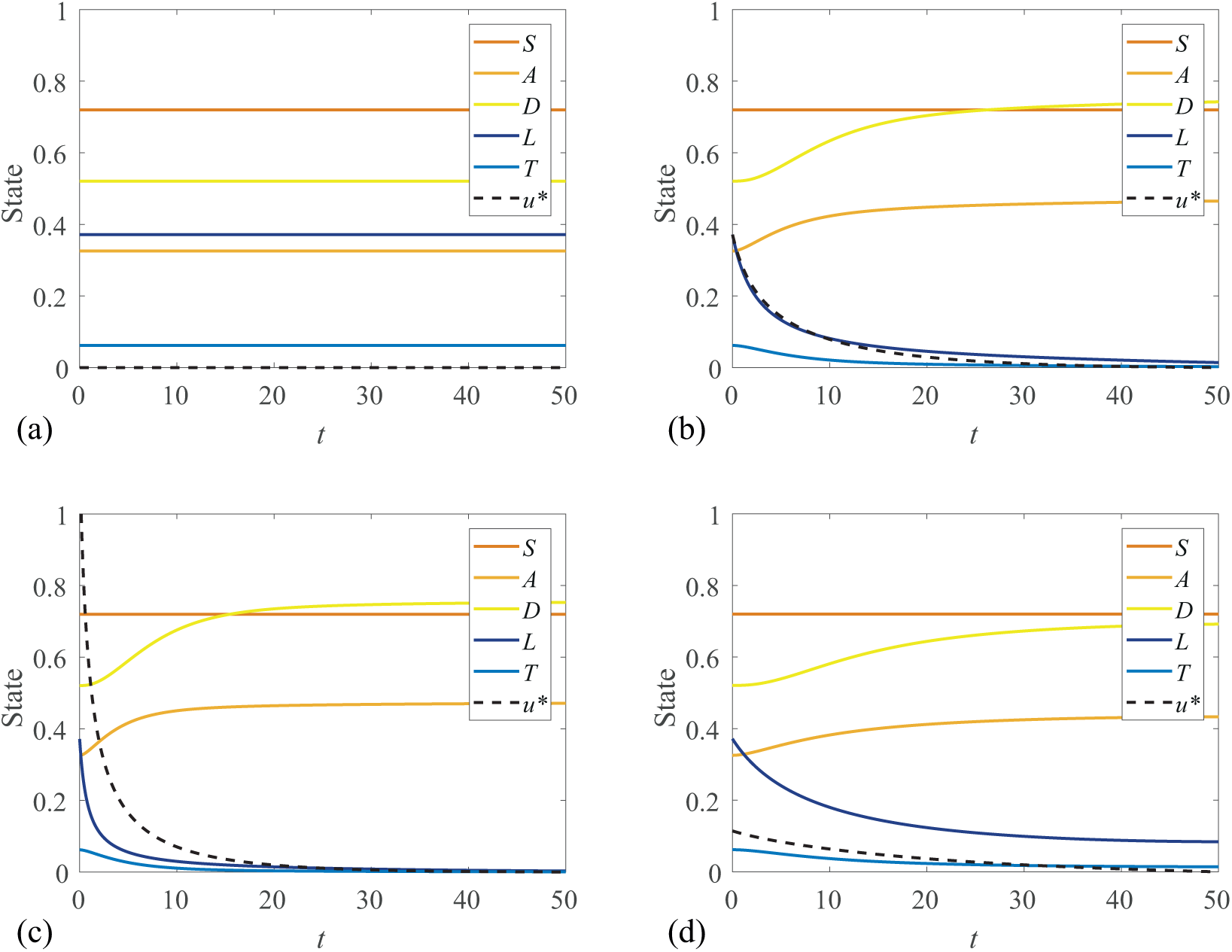
Application of a continuous optimal control (black dashed line) for various pay-off weightings *a*_1_ and *a*_2_. The corresponding pay-off, *J,* is also given. (a) Coexisting steady state solution with no control applied. (b) Equal weighting [*a*_1_, *a*_2_] = [1, 1], *J* = 0.7167. (c) Leukaemia weighted more heavily [*a*_1_, *a*_2_] = [0.1, 1], *J* = 0.2288. (d) Control weighted more heavily [*a*_1_, *a*_2_] = [1, 0.1], *J* = 0.2262. These figures are produced with immune response parameters *α* = 0.015, *γ* = 0.1.

For each of the optimal controls presented in Figure 5, we include an estimate of *J,* calculated by evaluating Equation (13) with the trapezoid rule. It is critical to note that these pay-offs should not be directly compared with each other. This kind of comparison would be meaningless as each result corresponds to different choices of *a*_1_ and *a*_2_, and these values explicitly contribute to *J*. For example; suppose an optimal control with pay-off weightings *a*_1_ and *a*_2_ is computed to have a pay-off of *J*_1_. Recomputing the optimal control with weightings 2*a*_1_ and 2*a*_2_ would produce a near identical optimal control and corresponding state, with slight deviation due to floating-point error. However, the corresponding pay-off *J*_2_ would be twice as large.

No pay-off is calculated for the uncontrolled steady state solution (Figure 5a) as the choice of *a*_1_ and *a*_2_ would be arbitrary. In this sense, computed pay-offs are not useful for comparing the outcome of *treatment* versus *no treatment* as there is no meaningful pay-off associated with no treatment. Rather, computed pay-offs can be used for comparison with other controls applied to a system with identical parameters to check whether or not they are comparable in outcome to the optimal control, noting that the response of the state will also change if the control changes.

To illustrate this point, we compare the optimal control obtained in Figure 5b to other potential treatment scenarios. In Figure 6 we compare four different dosing strategies where the same total amount of drug is applied using different temporal regimes. Our calculations of *J* provide a measure of how much the optimal result (Figure 6a) outperforms the other heuristically-determined dosing strategies. Applying the control at a constant rate for the full duration of the simulation (Figure 6b) produces a worse outcome than clinically-motivated cyclic treatment designs; applying the control at a greater level for a shorter duration in one (Figure 6c) or two (Figure 6d) cycles [5]. Due to the quadratic control term in Equation (13), despite applying the same total dosage, the control contributes more to the pay-off in Figure 6c than 6d, but this is outweighed by the benefit of reducing the leukaemic population more quickly. The optimal control framework provides us with tools to generate treatment hypotheses and assess the efficacy of different treatment protocols relative to one another and to the theoretical optimum for a given set of parameters.

**Fig. 6.**
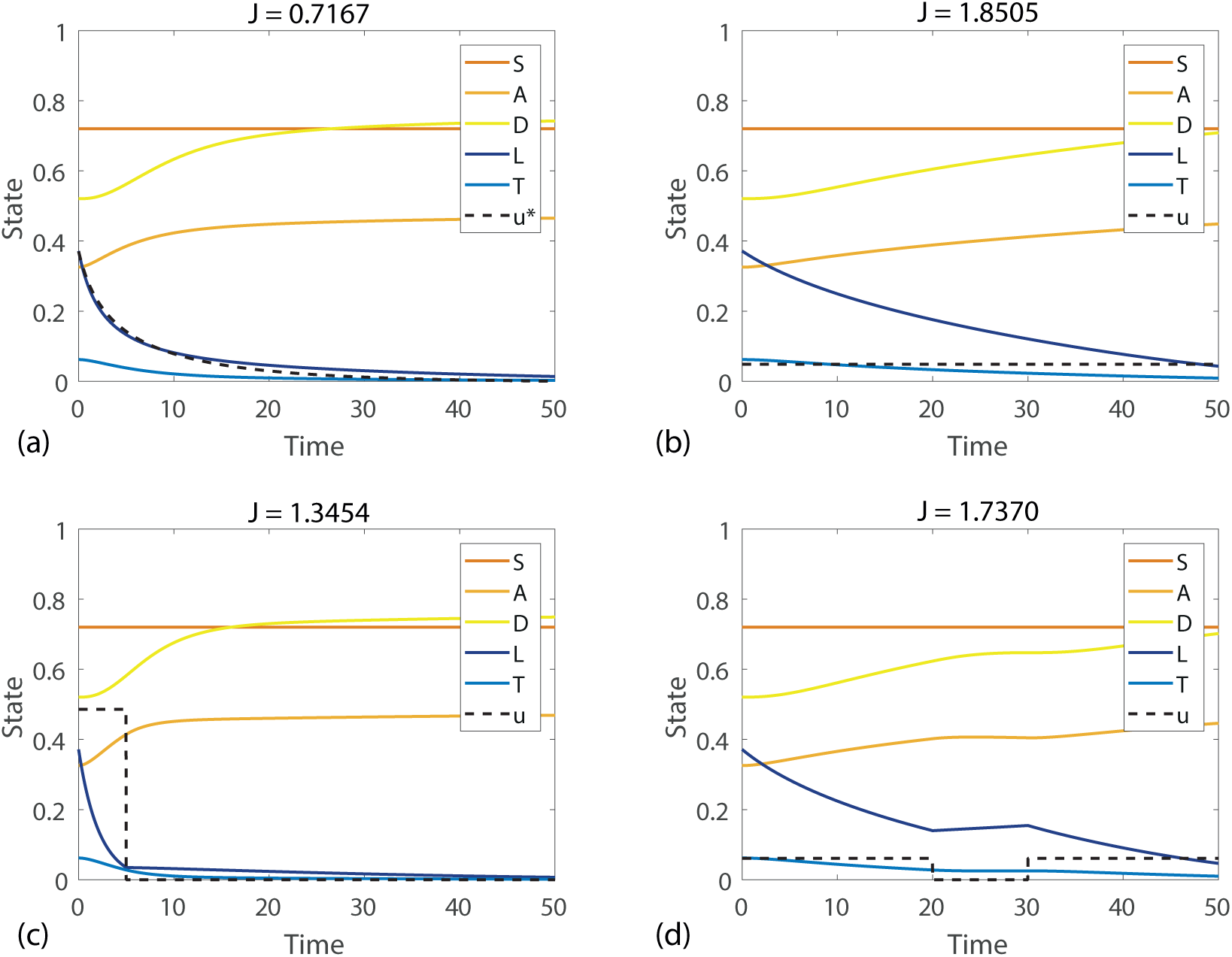
Comparison of (a) continuous optimal control with other possible controls of the same total dosage; (b) control applied at a constant rate over the entire duration, (c) control applied at a higher rate over a short cycle and (d) control applied in two cycles. These figures are produced with *a*_1_ = *a*_2_ = 1 and immune response parameters *α* = 0.015, *γ* = 0.1.

### 4.3 Bang-bang optimal control

In addition to considering continuous controls, it is also relevant to consider discontinuous bang-bang controls as this kind of *on-off* control could be thought to be more clinically relevant than a continuous setting. Bang-bang control problems require a specified bound on the control variable. A bang-bang optimal control takes the value of either the upper or lower bound with finitely many switching points over an interval. As a starting point we re-consider Equation (12) and note that a control will be either bang-bang optimal or singular if the pay-off function is linear in the control term. A pay-off that should produce a bang-bang or singular optimal control of Equation (12) is to minimise

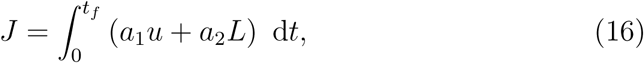

subject to *b*_1_ ≤ *u* ≤ *b*_2_. We can construct the Hamiltonian as *H* = ℒ+ **λ***f,* where ℒ is the integrand of Equation (16), **λ** = [λ_1_, λ_2_, λ_3_, λ_4_, λ_5_] and *f* is the right hand side of Equation (12), giving

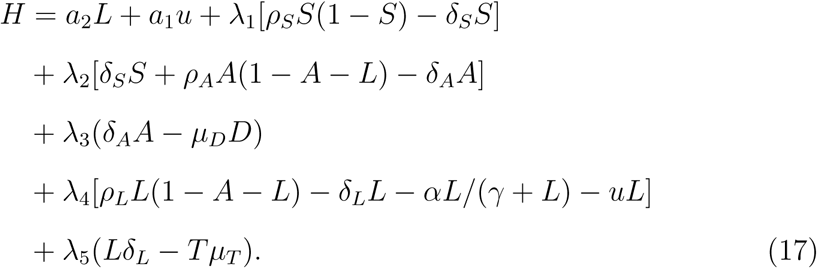

As for the continuous control case, we differentiate the Hamiltonian with respect to our control variable *u*. With a linear pay-off, however, the result no longer contains *u*. Rather than solving for *u*, we define a switching function, *ψ*(*t*), given by

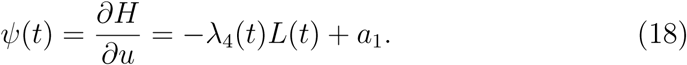

From PMP [51], it is implied that the Hamiltonian will be minimised under the following conditions,

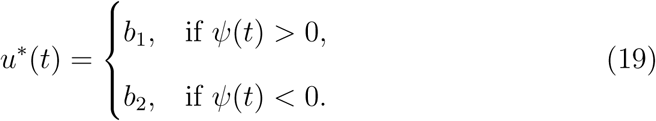

Conditions in Equation (19) produce a bang-bang control. Here, the control variable takes a value of either its upper or lower bound. Notably, Equation (19) omits the case where *ψ*(*t*) = 0, as a bang-bang optimal control requires that *ψ*(*t*) = 0 only at discrete points, if at all [14]. If *ψ*(*t*) = 0 for any finite interval aside from isolated points, the control is singular. Singular controls are most commonly encountered in cases where the Hamiltonian is linear in the control variable but non-linear in some state variables [12]. When *ψ*(*t*) = 0 over an interval, the Hamiltonian is not a function of the control, so the state and co-state variables no longer determine the control [12]; over this interval the control is determined by requiring *∂H/∂u* = 0. Our control problem defined by Equation (12) and Equation (16) is not singular, so we do not discuss singular controls further.

Our co-state equations for **λ** are found as *∂H/∂***x** = -d**λ***/*d*t*. The co-state in the bang-bang control problem is given by,

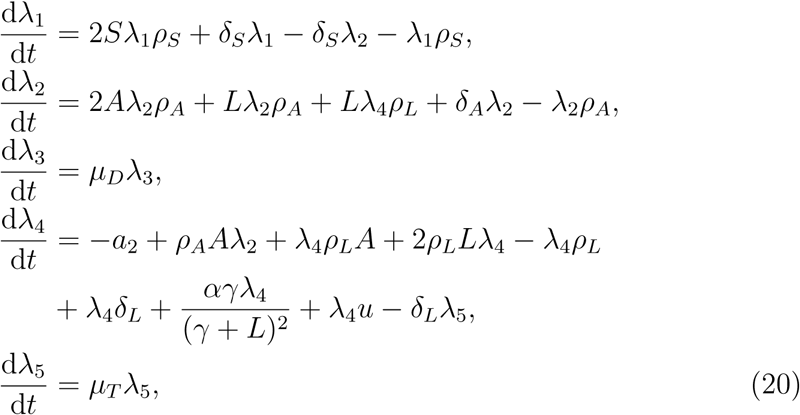

and we note that Equation (20) is subtly different to Equation (15), as the first term of the fourth line of Equation (20) is the constant -*a*_2_, and no longer depends on *L*.

The transversality condition, Equation (11), gives the final time conditions on the co-state, [λ_1_(*t*_*f*_), λ_2_(*t*_*f*_), λ_3_(*t*_*f*_), λ_4_(*t*_*f*_), λ_5_(*t*_*f*_)] = [0, 0, 0, 0, 0]. Assuming again that the initial state is known; [*S*(0), *A*(0), *D*(0), *L*(0), *T* (0)], it is now possible to determine the optimal bang-bang control and corresponding optimal state and co-state through a two-point BVP that we solve using the FBSM, as in the continuous control case. It is not necessary to modify the FBSM algorithm to find bang-bang optimal controls, though care must be taken in how the control is updated between iterations. This is discussed further in Section 4.4. Depending on the numerical scheme used to integrate the state and co-state equations through time, the discontinuous nature of the bang-bang control may require careful handling. Solutions are provided in Figure 7 for various weighting on the control parameters. In the continuous control case, when *a*_1_ > *a*_2_, placing a greater weighting on the negative impact of the control than the negative impact of the leukaemic stem cells; we observed that the control is applied at a lower level than when *a*_1_ < *a*_2_. The optimal bang-bang control must take either the upper or lower bound of the specified range. As such, in the bang-bang control case the pay-off weighting parameters determine not the level at which the control is applied, but rather the times at which the control switches from one bound to the other, hence the name *switching function* given to Equation (19).

**Fig. 7.**
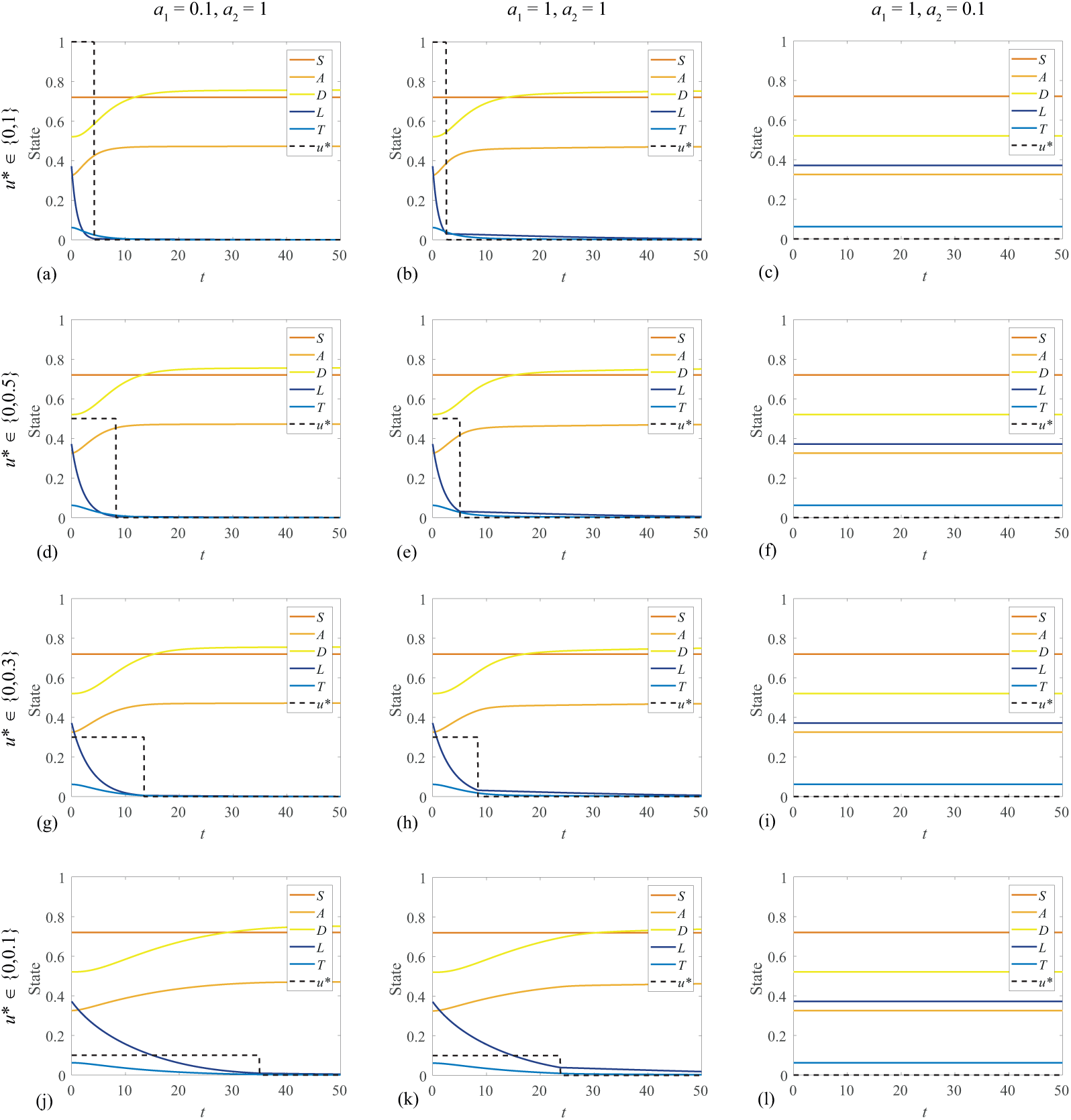
Bang-bang control solutions for various weightings on control and leukaemia in the pay-off (*a*_1_ and *a*_2_ respectively), with different control upper bounds. These figures are produced with immune response parameters *α* = 0.015, *γ* = 0.1.

In Figure 7 it is clear that when the upper bound on the control is higher, meaning in this context the maximum amount of chemotherapy that can be applied at any given time is higher, the control switches to the lower bound earlier. In this case the lower bound corresponds to *u* = 0, or no chemotherapy being applied (control *switched off)*, though this is not required of the method. The interaction between the control and state in Equation (20) means that the cumulative amount of control applied is not the same for different bounds on the control. In Figure 5 we demonstrate that for a continuous control with *a*_1_ = 1, *a*_2_ = 0.1, a small amount of control is applied. For the bang-bang case with the same weighting, we observe in the rightmost column of Figure 7 that for a range of control upper bounds, the control is not switched on at all - implying that with such a pay-off, it is optimal not to apply the control. One may suppose that for a sufficiently small upper bound that the control would turn on even with this pay-off, however a lower upper bound on the control also reduces the impact the control has on the state.

Due to the immune response incorporated in Section 3, a sufficiently small leukaemic population will tend towards extinction rather than grow back to a coexisting steady state. Because of this, we observe in Figure 7 that the control switches off before the leukaemic stem cells are totally eradicated - the immune response is sufficient once the leukaemic population is sufficiently low. This is most evident in Figure 7k, where we can see that the population of leukaemic stem cells is declining but has not become extinct by the final time, *t* = 50. In absence of the immune response incorporated in Section 3, we would observe the leukaemic population increasing as soon as the control is switched off, since the healthy steady state would be unstable; applying fixed final time bang-bang optimal control to the original model produces outcomes that are mathematically optimal but physically undesirable.

In our discussion of continuous controls, we note the fixed final time as a limitation, since changing the final time can change the profile of the optimal control and state. In general the same is true of bang-bang controls with fixed final times, though in some instances that we consider the optimal bang-bang control does not change significantly if the final time is changed. For example; the optimal switching times and corresponding optimal states in the leftmost column of Figure 7 do not change significantly if the final time is increased to *t* = 100, because by *t* = 50 we see that *L ≈* 0 and *u* = 0, so neither contributes significantly to the pay-off in the interval 50 < *t* ≤ 100. For these cases the control is not costly relative to the leukaemia (*a*_1_ < *a*_2_) so it is applied at the upper bound until the leukaemic stem cell population is virtually eradicated before switching off.

For this particular system, we only obtain bang-bang optimal controls with a single switching time. We are able to verify these bang-bang optimal controls through an exhaustive search of all possible bang-bang controls by specifying the switching time, directly calculating the pay-off and determining the switching time that minimises the pay-off. For all cases considered in Figure 7 the switching time identified via exhaustive search is in agreement. It is also possible that the optimal bang-bang control may switch between the upper and lower bounds numerous times, producing multiple ‘bangs’. Bang-bang optimal controls that exhibit multiple bangs can be identified using the FBSM without modification, though it is more difficult to find a convergent bang-bang optimal control with multiple bangs. Similarly, without knowing a priori how many switching times to expect, an exhaustive search for multiple bangs is not computationally feasible.

### 4.4 Convergence and control updating

In this section we examine the convergence behaviour of solutions to the optimal control problems presented in this work. Convergence behaviour of numerical solutions to optimal control problems is influenced by multiple factors. In particular, we discuss the initial guess of the control, convergence criteria, control updating and pay-off weightings. These factors influence not only the number of iterations required to reach a converged numerical solution, but also whether or not a converged solution will be reached at all.

Holding all other factors constant, provided that the initial guess for the control is sensible, the initial guess does not have a significant impact on whether or not a converged result is reached for the control problems considered in this work. However, convergence is typically reached with fewer iterations when the initial guess is relatively closer to the true value of the optimal control. For simplicity we use the initial guess *u* ≡ 0 for all results presented in this work, while acknowledging that more thoughtful choices may deliver convergence in fewer iterations.

For optimal control results presented in the previous sections, we determine whether convergence has been achieved after each iteration based on the relative difference between the updated control, *u*_updated_, and the old control, *u*_old_. If this relative difference is sufficiently small, the updated control is accepted as the optimal control. A typical relative difference convergence criterion requires

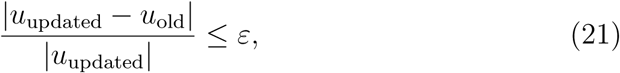

where 0 < *ε* ≪ 1 is the desired relative tolerance. Following [37], we adjust Equation (21) to allow for a control of the form *u* ≡ 0, giving

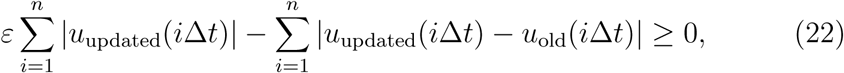

where *t* = *i*Δ*t*, Δ*t* is the numerical time step and *n* is the number of nodes in the time discretisation. The absolute value is taken to ensure that positive differences are not offset by negative differences that could otherwise result in incorrectly detecting convergence. The choice of convergence criterion and acceptable tolerance depends on the particular problem at hand, and may need to be adjusted to be appropriate for another control problem. In some instances, it may be necessary to check convergence of the state and co-state as well as the control, particularly if the state response to control is sensitive. For the control problems studied in this work, we find that state and costate respond predictably to the control, and convergence of the control is accompanied by convergence of that state and co-state. As such we do not explicitly check for convergence of the state and co-state.

In each iteration of the FBSM we recalculate the control, *u*_new_, based on the newly calculated state and co-state solutions and associated optimality criterion, as discussed in Section 4.2 for the continuous control and Section 4.3 for the bang-bang control. Typically, *u*_new_ is not used directly as the control for the next iteration of the FBSM, but rather we form an updated control *u*_updated_ as a weighted combination of *u*_new_ and the control from the previous iteration, *u*_old_. The motivation for this is two-fold; first, an appropriately weighted control updating scheme can speed up convergence; and second, for many optimal control problems, a direct update of *u*_updated_ = *u*_new_ will fail to produce converging results at all. A common approach is to update the control based on a convex combination, such that the total weightings sum to one, of the new and previous control(s). In this work we use a constant linear weighting, with 0 < *ω* < 1, giving

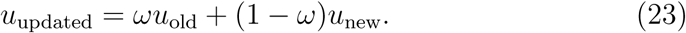

We find that the best choice for *ω* depends not only on the form of the control, continuous or bang-bang, but also on model parameters such as the pay-off weightings. There is a trade-off between the number of iterations required to obtain convergence, and actually converging at all; a larger *ω* typically is more likely to produce converging solutions, but this also means that the control changes less each iteration, so more iterations are required. For example, a weighting of *ω* = 0.7 was sufficiently large that all continuous control solutions presented in Figure 5 converged to a relative tolerance of *ε* = 1 *×* 10^−3^. For *ω* = 0.6 only Figure 5d converges, and for *ω* = 0.8, all solutions in Figure 5 converge but require more iterations than when *ω* = 0.7.

Convergence in the bang-bang control case typically requires larger *ω* and more iterations than the continuous controls. In the rightmost column of Figure 7, there is no concept of convergence as the control never switches on. Only Figure 7j and Figure 7k converge to a relative tolerance of 1*×*10^−3^ for *ω* = 0.7, with *ω* = 0.9 being sufficient for convergence of all remaining solutions aside from Figure 7b, where we set *ω* = 0.95.

It is clear that the best control updating scheme depends on the particular problem; and a scheme that works well for one problem may not necessarily work at all for another. When solving control problems, it may be necessary to try a range of updating schemes to achieve convergence. In this work we only consider constant weighted updating, though there are more sophisticated updating schemes that shift the weighting towards *u*_new_ as the number of iterations increase [37]. In Figure 8 we examine the influence of the control update weighting *ω*, and the pay-off weightings, *a*_1_ and *a*_2_, on the convergence behaviour of the bang-bang control problem studied in Section 4.3. Specifically, we consider the case where 0 ≤ *u* ≤ 0.5, and determine that a solution has converged if it meets a relative tolerance of *ε* = 1 *×* 10^−3^ within 250 iterations. In each panel of Figure 8 we observe three *regions*: in region I we have no concept of convergence as the control never switches on; in region II we find that the optimal control problem does not converge; and in region III we observe convergence. Not all simulations conform strictly to these regions since the boundary between the different regions is not always sharp and welldefined. However, broadly speaking, these three regions capture the essence of the convergence behaviour that we observe. These regions are constructed based on discrete simulations of the problem for 0 < *a*_1_ ≤ 10 and 0 ≤ *a*_2_ ≤ 10, each in increments of 0.1. The case where *a*_1_ = 0 is excluded as this corresponds to no cost associated with applying the control, so there is no sense of convergence.

**Fig. 8.**
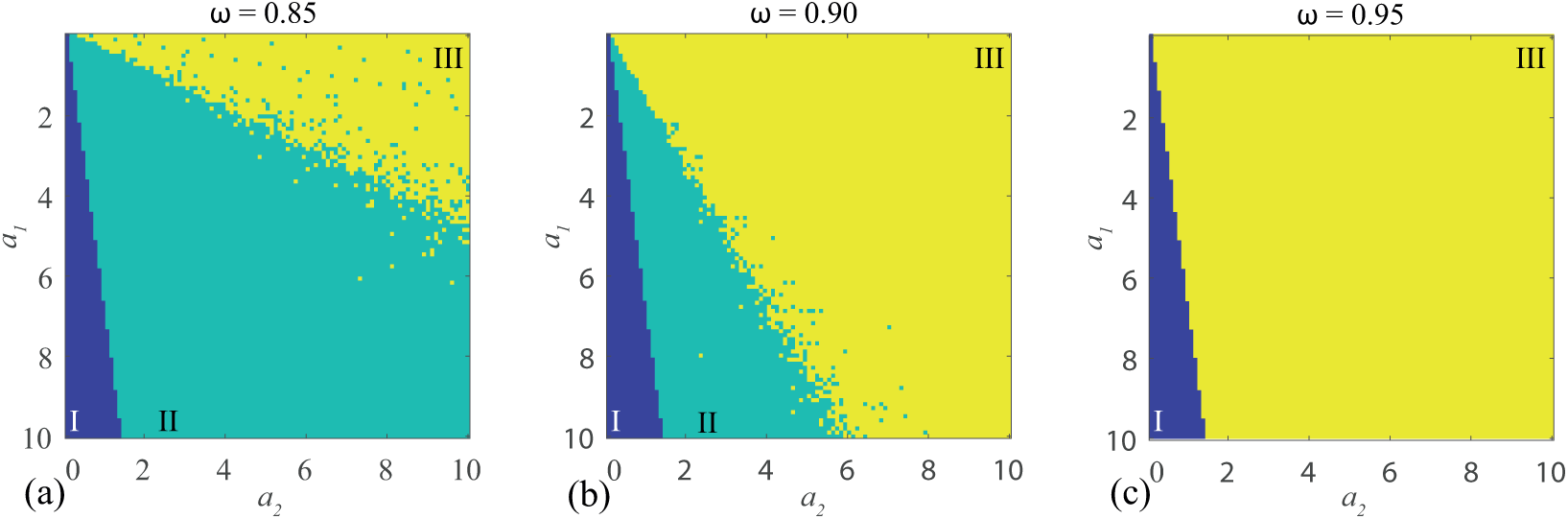
Convergence behaviour for (a) *ω* = 0.85, (b) *ω* = 0.9, and (c) *ω* = 0.95, with *a*_1_ and *a*_2_ ranging from 0 to 10 in increments of 0.1, excluding *a*_1_ = 0. In region I (dark blue) we have no concept of convergence as the control never switches on. In region II (light blue) we find that the optimal control problem does not converge, and in region III (yellow) we observe convergence. These figures are produced with immune response parameters *α* = 0.015, *γ* = 0.1.

From Figure 8 it is clear that convergence is achieved in a larger region of the (*a*_1_, *a*_2_) parameter space when *ω* is increased. However, it is important to note that achieving convergence in this context only implies that Equation (22) is satisfied, and does not necessarily mean that a suitable bang-bang control is obtained. While some controls corresponding to individual simulations in Figure 8c are suitable bang-bang controls; a portion are approaching bang-bang but require additional iterations to accurately calculate the control around the switching point. The weighting applied in Equation (23) has the effect of smoothing *u* during intermediate iterations of the FBSM; this smoothness is gradually reduced as the control converges to the optimal switching point. Since *ω* explicitly influences the relative amount that the control can differ between iterations, if a larger *ω* is required to achieve convergence for a given problem, it may also be necessary to reduce the convergence tolerance *ε* to ensure that the resulting control is sufficiently bang-bang.

## 5 Conclusion and Outlook

In this work we consider a haematopoietic stem cell model of AML that incorporates competition between leukaemic stem cells and blood progenitor cells within the bone marrow niche. We incorporate a biologically appropriate immune response in the form of a Michaelis-Menten term. This modification is mathematically convenient because of the impact it has on the steady states, and biologically relevant because the immune response is known to play an important role in cancer progression and treatment. With a view to identifying the optimal way to apply a treatment such as chemotherapy to the model, we formulate and solve optimal control problems corresponding to multiple objectives and constraints. This includes quadratic pay-off functions, yielding continuous controls, as well as linear pay-off functions, yielding discontinuous bang-bang controls.

We provide a brief overview of optimal control theory, with a focus on the necessary conditions derived from Pontryagin’s Maximum Principle. This approach formulates the optimal control problem as a coupled multi-species two-point boundary value problem. The resulting optimal control problem is solved numerically using the iterative FBSM. The algorithm for the FBSM is discussed, with a focus on highlighting typical issues that may arise in implementing optimal control. Suggestions are provided for overcoming these issues. In particular, we focus on factors that influence the convergence of the FBSM; not only in terms of the number of iterations required, but also whether it converges at all. These factors include the initial guess for the control, the convergence criterion, the method of updating the control, the associated weighting placed on controls from prior iterations and parameters such as pay-off weightings, and in the bang-bang control case, the control bounds.

For the model we consider; a well informed initial guess for the control may reduce the number of iterations required for convergence, but any sensible guess should not prevent convergence. Most critically, we show that the method of updating the control and the associated weight placed on the control from the previous iteration has a significant impact on whether or not convergence will be achieved, as do the weights in the pay-off function. In the bang-bang control case, we observe that increasing the upper bound on the control can prevent convergence, holding all other factors constant; in this case, placing a greater weight on the solution from the previous iteration may produce convergence.

There are many potential avenues to extend the ideas explored in this work. Here, we have incorporated the control via a simple mechanism, and more sophisticated pharmacokinetic processes such as drug absorption and metabolism could be incorporated to increase the biological detail captured by the model, but this additional biological detail comes at the cost of increasing the number of unknown, and possibly unmeasurable parameters. Therefore, care must be exercised in following up this kind of extension. In the main document we consider the most fundamental case of a control that only impacts leukaemic cells, however the methodology extends to a control affecting multiple species. We demonstrate this extension in the supplementary material. The control problems presented in this work could be reformulated as fixed final state problems, leaving the final time free to vary which could be more clinically relevant than specifying the final time. With the introduction of an immune mechanism to the model, it is also possible to consider a control based around immunotherapy.

A recent idea of great interest in clinical cancer research is the possibility of introducing an interval of time during treatment in which no chemotherapy is applied. This kind of intervention is reminiscent of a bang-bang control, and is often referred to as a *drug holiday* [60]. There is some evidence to suggest that drug resistance of tumour cells may reduce with time so that patients experience an improved response to chemotherapy following a drug holiday [33,34,55]. This application of a drug in an on-off fashion parallels the idea of the bang-bang controls we consider in this work and so it would be interesting to formulate the concept of designing a drug holiday in terms of a bang-bang optimal control problem by extending the model to include acquired drug resistance and using the algorithms and concepts demonstrated in this work.

## Acknowledgments

This work is supported by the United States Air Force Office of Scientific Research (BAA-AFRL-AFOSR-2016-0007) and the Australian Research Council (DP170100474). Computational resources were provided by the eResearch Office at QUT. We appreciate the helpful referee comments.

