## Supplementary Information for "Optimal control of acute myeloid leukaemia"

**Contents**

1 Arbitrary initial conditions 2

2 Control affects all proliferative cells 4

References 7

* Corresponding author

### 1 Arbitrary initial conditions

Optimal control results in the Main paper are produced with an initial condition corresponding to a stable steady state of the system. In Figure 1 we present results demonstrating that the optimal control techniques do not require such an initial condition. To produce these results we choose an arbitrary initial condition, solve the system numerically up to some transient time without control, $u \equiv 0$, then apply the FBSM from that transient time to the final time to determine the optimal control.

These results adhere to our intuitive expectations; with a larger initial leukaemic population, a greater amount of control is applied, for example compare Figure 1d and Figure 1e. Similarly, if there is a larger initial haematopoietic stem cell population, a smaller amount of control is required, for example compare Figure 1a and Figure 1c.

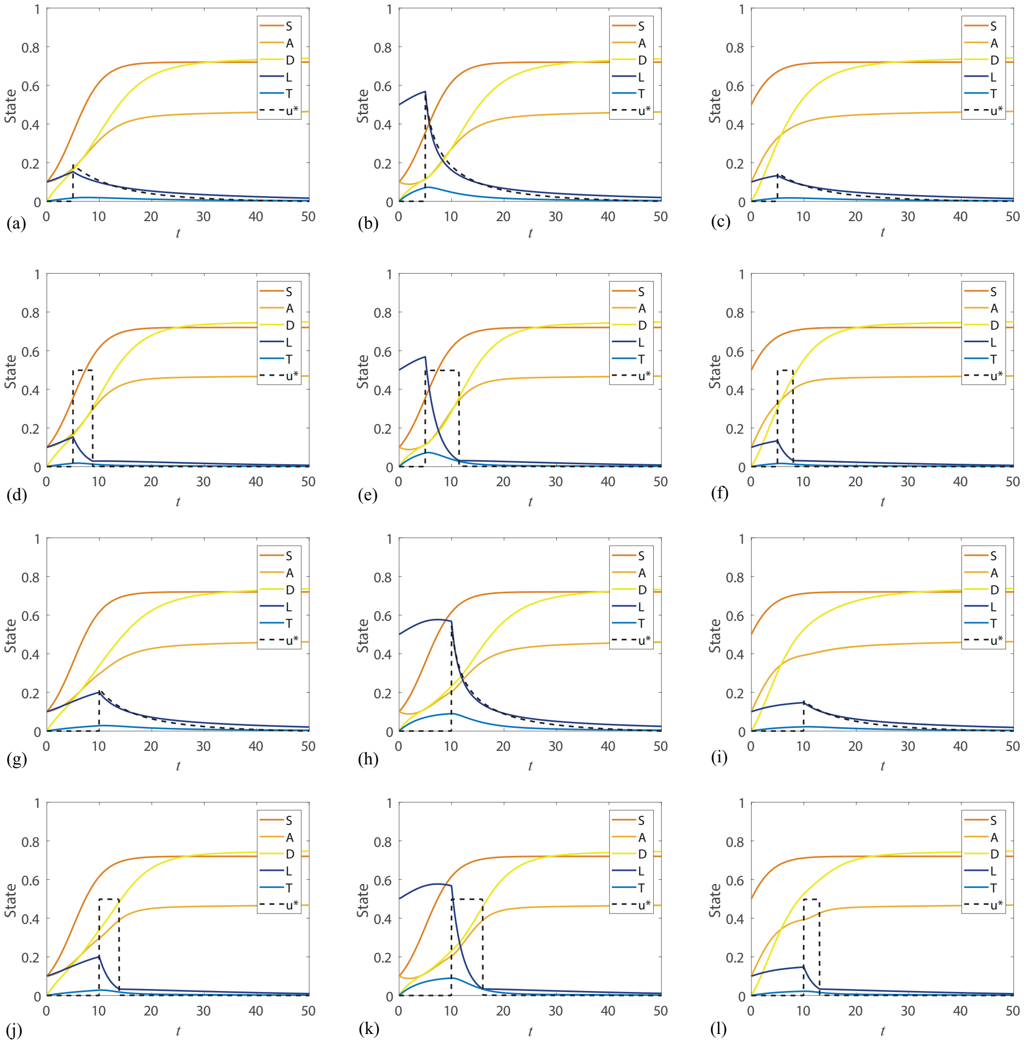


Fig. 1. Results are produced for a range of arbitrary initial conditions subject to a transient growth phase prior to application of a control. Columns from left to right correspond to initial conditions of $[0.1, 0.1, 0, 0.1, 0]$, $[0.1, 0.1, 0, 0.5, 0]$ and $[0.5, 0.1, 0, 0.1, 0]$. Control is applied from $t = 5$ in Figures (a-f) and from $t = 10$ in Figures (g-l). The first and third rows correspond to the continuous control case, while the second and fourth rows show the corresponding bang-bang case with an upper control bound of $u = 0.5$. For all results in this figure, $a_1 = a_2 = 1$.

#### 2 Control affects all proliferative cells

In reality, typical chemotherapy treatment of AML uses cytotoxic drugs that affect not only leukaemic cells but all proliferative cells [1]. In this section we demonstrate that our analysis can be extended to consider the case where the control also impacts the proliferative healthy cells; $S$ and $A$. We extend our state equations (Equation 12, Main paper) by introducing terms with parameters $r_1$ and $r_2$ describing the rates at which the control, $u$, affects the $S$ and $A$ populations relative to its effect on the $L$ population:

$$\begin{aligned}
\frac{\mathrm{d}S}{\mathrm{d}t} &= \rho_S S(K_1 - Z_1) - \delta_S S - r_1 u S, \\
\frac{\mathrm{d}A}{\mathrm{d}t} &= \delta_S S + \rho_A A(K_2 - Z_2) - \delta_A A - r_2 u A, \\
\frac{\mathrm{d}D}{\mathrm{d}t} &= \delta_A A - \mu_D D, \\
\frac{\mathrm{d}L}{\mathrm{d}t} &= \rho_L L(K_2 - Z_2) - \delta_L L - \frac{\alpha L}{\gamma + L} - uL, \\
\frac{\mathrm{d}T}{\mathrm{d}t} &= \delta_L L - \mu_T T. \qquad (1)
\end{aligned}$$

We define a pay-off function to minimise, such as

$$J = \int_0^{t_f} \left(a_1 u^2 + a_2 L^2\right)\, \mathrm{d}t, \qquad (2)$$

for the continuous control case, or

$$J = \int_0^{t_f} (a_1 u + a_2 L)\, \mathrm{d}t, \qquad (3)$$

for the bang-bang control case. Recalling that $a_1$ weights the relative importance placed on the negative impact of the control, we could at this point consider reducing $a_1$ relative to $a_2$, as we have now explicitly accounted for the direct negative effect of the control on the healthy cells. We could also explicitly include $S$ and $A$ in the pay-off. The motivation for this becomes apparent when considering a control that affects $S$ and $A$ more than $L$; $r_1, r_2 > 1$, as in Figure 2g-l, as we see the healthy cell counts become very low during the treatment period, which may be clinically undesirable. These terms would need to be weighted negatively, as the pay-off is a quantity to be minimised.

Following the procedure in the Main paper we construct the Hamiltonian and find the optimal control by setting $\partial H/\partial u = 0$, or define a switching function in the bang-bang control case. Co-state equations for $\boldsymbol{\lambda}$ are found by setting $\mathrm{d}\boldsymbol{\lambda}/\mathrm{d}t = -\partial H/\partial \mathbf{x}$, giving

$$\begin{aligned}
\frac{\mathrm{d}\lambda_1}{\mathrm{d}t} &= 2S\lambda_1\rho_S + \delta_S\lambda_1 - \delta_S\lambda_2 - \lambda_1\rho_S + \lambda_1 r_1 u,\\
\frac{\mathrm{d}\lambda_2}{\mathrm{d}t} &= 2A\lambda_2\rho_A + L\lambda_2\rho_A + L\lambda_4\rho_L + \delta_A\lambda_2 - \delta_A\lambda_3 - \lambda_2\rho_A + \lambda_2 r_2 u,\\
\frac{\mathrm{d}\lambda_3}{\mathrm{d}t} &= \mu_D\lambda_3,\\
\frac{\mathrm{d}\lambda_4}{\mathrm{d}t} &= -2a_2L + \rho_A A\lambda_2 + \lambda_4\rho_L A + 2\rho_L L\lambda_4 - \lambda_4\rho_L\\
&\quad + \lambda_4\delta_L + \frac{\alpha\gamma\lambda_4}{(\gamma+L)^2} + \lambda_4 u - \delta_L\lambda_5,\\
\frac{\mathrm{d}\lambda_5}{\mathrm{d}t} &= \mu_T\lambda_5,
\end{aligned} \tag{4}$$

in the continuous control case. The corresponding co-state equations for the bang-bang control case are subtly different to Equation (4), as the first term of the fourth line of Equation (4) is the constant $-a_2$, and no longer depends on $L$. The transversality condition again provides the final time conditions on the co-state, $[\lambda_1(t_f), \lambda_2(t_f), \lambda_3(t_f), \lambda_4(t_f), \lambda_5(t_f)] = [0, 0, 0, 0, 0]$. Given an initial state; $[S(0), A(0), D(0), L(0), T(0)]$, we can solve the resulting two-point BVP using the FBSM, as detailed in the Main paper. Results are provided in Figure 2 corresponding to various parameter combinations.

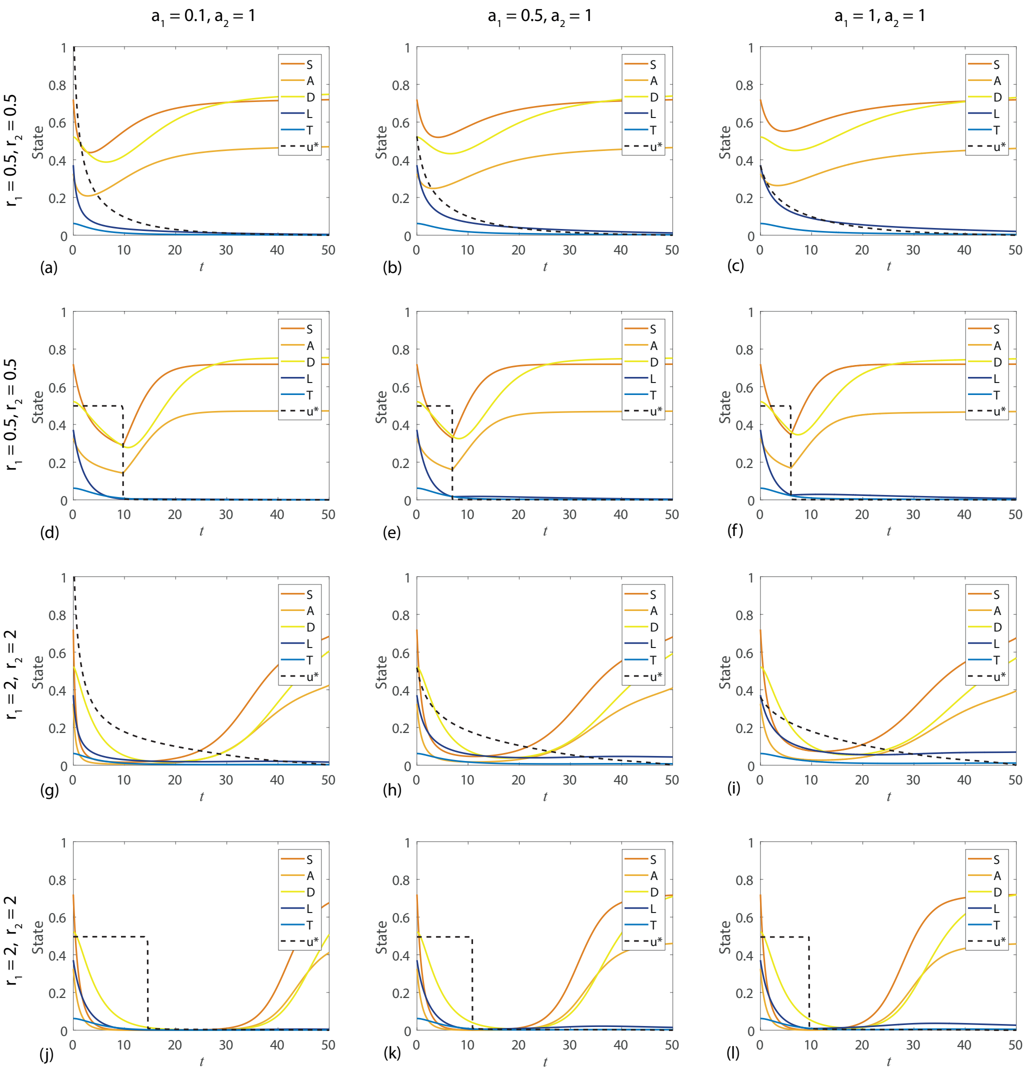


Fig. 2. Results are produced for a variety of parameters in the pay-off and rates that the control affects healthy cells. The continuous control case is presented in the first and third rows, while the second and fourth rows show the corresponding bang-bang case with an upper control bound of $u = 0.5$.
